# SignalGen: A Protein Language Model Based AI Agent For Optimal Signal Peptide Prediction

**DOI:** 10.1101/2025.11.12.688111

**Authors:** Joe McKenzie, Hung N. Do, Apoorv Shanker, Jessica Z. Kubicek-Sutherland, S. Gnanakaran

## Abstract

Signal peptides are short amino acid sequences attached to the N-termini of mature proteins. They play determinant roles in protein expression as well as localization of mature proteins. While sequence-based machine learning (ML) models have been developed to identify the signal peptide sequences given the full or mature protein sequences, no model has been created to design optimal signal peptides with the localization of the mature proteins taken into account. Here, we develop a ML model that considers the mature protein sequence, organism, and localization as inputs, encodes and processes them through a Latent Residual Transformer (LRT), and outputs the optimal signal peptide sequences for enhanced expression of the mature proteins, regardless of whether the proteins are non-native to the organism or de novo. The model is trained using the latest data from the UniProt database up until July 2025. Benchmarking of our ML model shows good performance in predicting the signal peptides for both human and non-human proteins from the UniProt database. Furthermore, our ML model is implemented with an artificial intelligence (AI) agent to enhance accessibility for the general scientific community. Findings from this study provide a framework for predicting optimal signal peptides for non-native protein expression of viral and bacterial vaccine candidates in human cells and for enhanced expression of *de novo* proteins.

## Introduction

Signal peptides are short amino acid sequences of typically 10 to 30 amino acids located at the N-termini of proteins^1,2^. Signal peptides play determinant roles in protein expression and the direction of processed proteins to target cellular compartments to carry out their functions^3^. Consequently, signal peptides act very much like zip codes for proteins and play a crucial role in the delivery of vaccine candidates in human cells^4^. In particular, signal peptides work by ensuring that antigens from a vaccine platform, such as mRNA, get expressed in the desired locations (such as intracellular, transmembrane, or extracellular) to trigger an immune response^5^. However, the detailed mechanisms upon which signal peptides are recognized to express and localize the mature proteins remain underexplored^6^. As a result, it remains difficult to design optimal signal peptides for protein expression, especially for viral and bacterial antigens in human cells^7^. Currently, most approaches rely on randomly testing a few *ad hoc* signal peptides to express viral and bacterial antigens in human cells to serve as vaccine candidates^8,9^. Therefore, the need arises for a rational design approach to improve the efficiency of non-native protein expression in human cells for vaccine design, especially for emerging pathogens.

Several computational tools have been developed to locate the signal peptide cleavage position in the immature protein sequence^10–13^. The SignalP team has contributed significantly to creating these types of tools^14^. SignalP 6.0 uses protein language models and is one of the current state-of-the-art signal peptide identification tools ^13^. Phobius is another popular signal peptide identification tool, but it was designed for distinguishing signal peptides and transmembrane α-helical regions^10,15,16^. Both signal peptides and transmembrane α-helical regions share large hydrophobic segments as characteristic features^1^. These tools have been considerable advances in the realm of signal peptide prediction. However, few validated tools have been developed for the task of signal peptide generation^17^. In fact, the concept of generating whole signal peptide sequences is relatively new^14^. SecretoGen is one of the most recent signal peptide generation tools that predicts a signal peptide for secretion of a mature protein sequence in the specified organism^18^. Nevertheless, there is currently no tools to generate a signal peptide sequence that will express a given mature protein in a desired cellular organelle. In a parallel study, we developed a structure-based optimization strategy to redesign the native signal peptides for enhanced protein expressions in human cells^19^. However, the strategy cannot be applied for *de novo* proteins, where no information of the native SPs is available.

Recent advances in the realm of AI and large language models (LLMs) have enabled the use of AI agents that can logically reason through complex tasks^20^. These agents are particularly useful in automating challenging and laborious tasks^21^. In this work, we implemented an AI agent for prediction of optimal signal peptide sequences for expression and target localization of both existing and *de novo* mature proteins in a given organism. We built a Latent Residual Transformer (LRT) model to predict the optimal signal peptide for the mature protein, organism, and specified subcellular location. This model is incorporated an AI agent to increase the accessibility to the broader scientific community. We provided two different ML models in predicting signal peptides, one of which was trained solely on human proteins, and the other was trained on all organisms. Our models introduce a new and efficient method for designing signal peptides to enhance expressions of native, non-native and *de novo* proteins into the desired compartments of a given organism.

## Results

We collected and reviewed protein data that contained the following information: mature protein sequences, signal peptide amino acid sequences, experimentally verified subcellular location, and associated organism from the UniProt Knowledgebase (https://www.uniprot.org/uniprotkb) up until July 2025 . Afterwards, we trained two different ML models, one solely on the 5,699 human protein data and one on the 55,620 proteins from all organisms within UniProt. The distributions of the subcellular localizations for the human and all-organism proteins, as well as the distributions of organisms in the all-organism dataset, can be found in **Figure 1**. The majority of proteins in both the human (32.7%) and all-organism (41.6%) datasets were secreted proteins, followed by proteins anchored in the cell membrane and the endoplasmic reticulum (**Figure 1**). However, more proteins were found in the periplasm than endoplasmic reticulum for all-organism proteins (**Figure 1B**). Beyond these cellular localizations, protein localization at other cellular organelles (total 57) occurred only at low frequencies and were grouped as “Others”. The distributions of the lengths of the signal peptides and mature proteins and the amino acid compositions in the signal peptide can be found in in **Supplementary Figures 1 and 2**. In both datasets, the signal peptides were typically less than 40 amino acids long and comprised of mostly leucines and alanines. The mature proteins were generally under 2,000 amino acids long in both the human-only and all-organism datasets.

**Figure 1.**
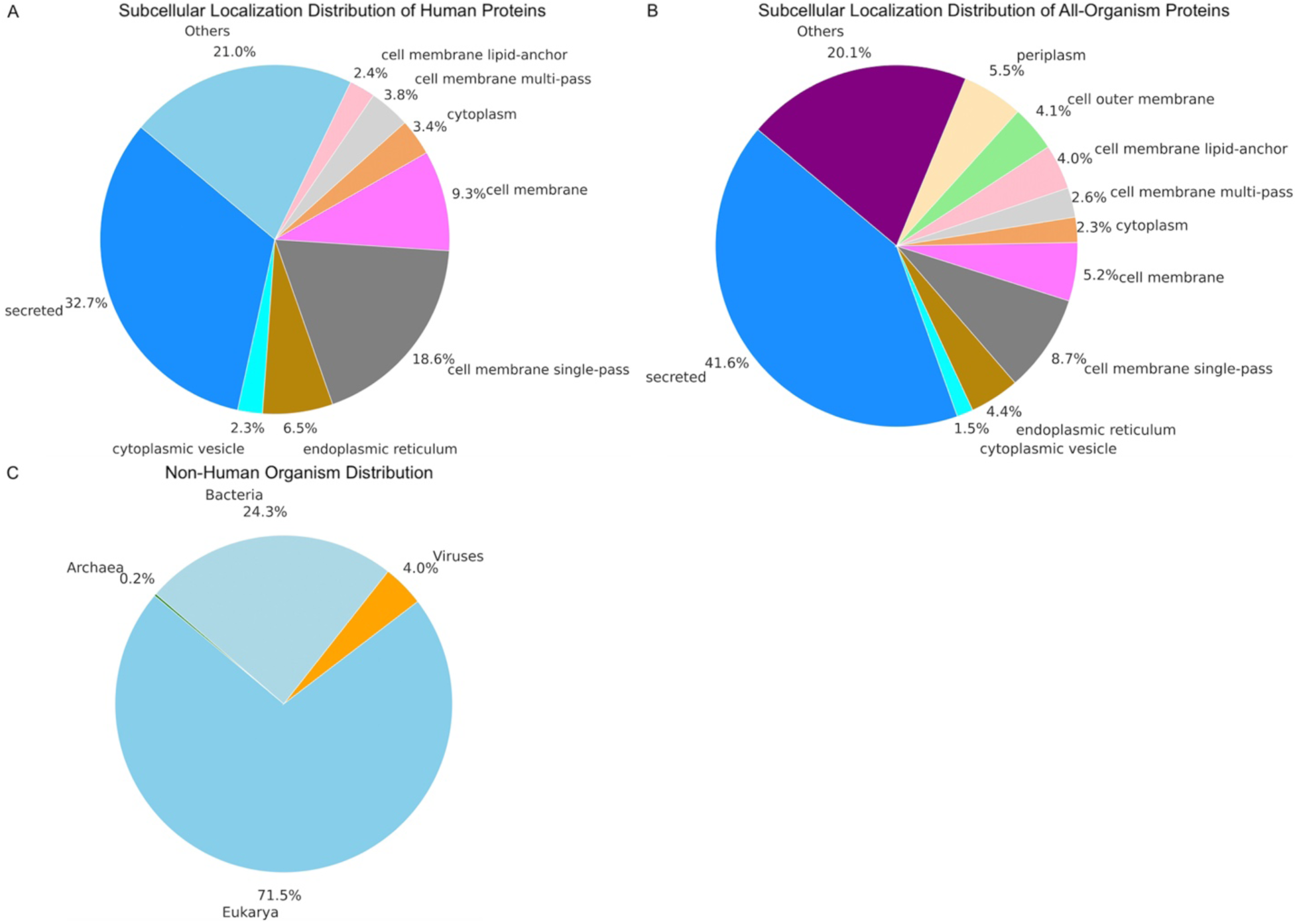
Distributions of auxiliary labels in the human-only and all-organism protein datasets. **(A)** Percentages of the 11 most common subcellular localizations in the human-only dataset with the other 57 grouped under the ‘Other’ category. **(B)** Percentages of the 11 most common subcellular localizations in the all-organism dataset with the other 57 grouped under the ‘Other’ category. **(C)** Percentages of non-human organisms in the all-organism dataset.

A detailed illustration of the training of our ML models is shown in **Supplementary Figure 3**. For training both the human-only and all-organism ML models, each dataset was randomly split into training and testing datasets using a ratio of 80:20 (**Figures 2**-**3** and **Supplementary Data 1**). The distributions of the subcellular localizations for the human and all-organism proteins in the training and test sets are shown in **Figure 3**. Here, we stratified each dataset so that the distribution of the localizations in the training and testing datasets of human proteins and of all-organism proteins would be similar (**Figure 3**). Furthermore, we confirmed that no data leakage or overlap took place between the training and test sets for training and evaluating both human-only and all-organism ML models (see **Methods** section). We achieved this goal by grouping each entry with the same signal peptide sequence together before splitting the data into training and testing sets.

**Figure 2.**
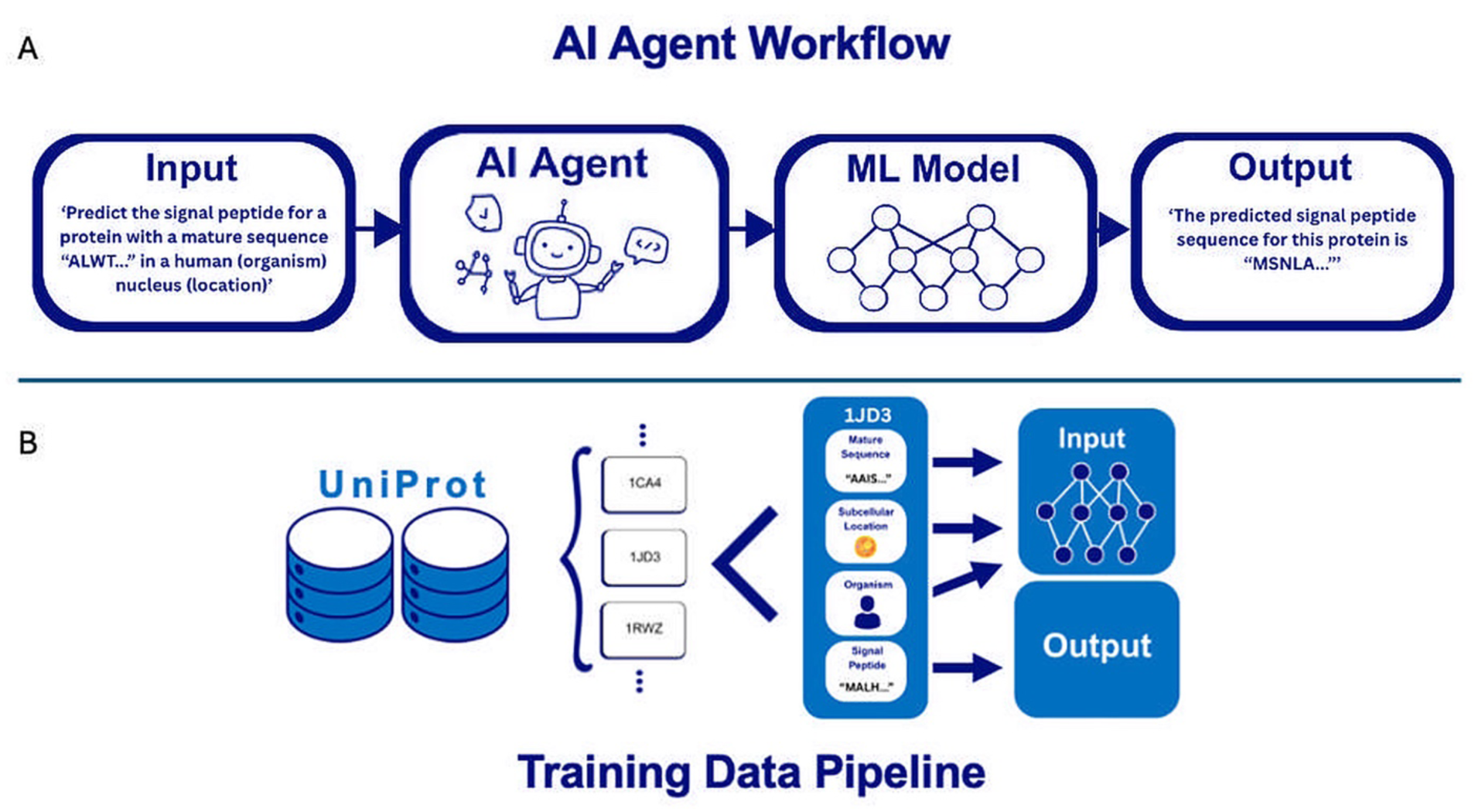
Schemes of the inference in deployment and preparation of training data. **(A)** The series demonstrates how a prompt is given to an AI agent, which then calls our trained ML model to predict the signal peptide sequences and output predictions to users. **(B)** The diagram shows how proteins are collected from a database and how information from collected proteins is used for either inputs or outputs for training the ML models. The signal peptide is the output, which is generated by the ML model learning from the mature protein sequence, the subcellular localization, and the organism.

**Figure 3.**
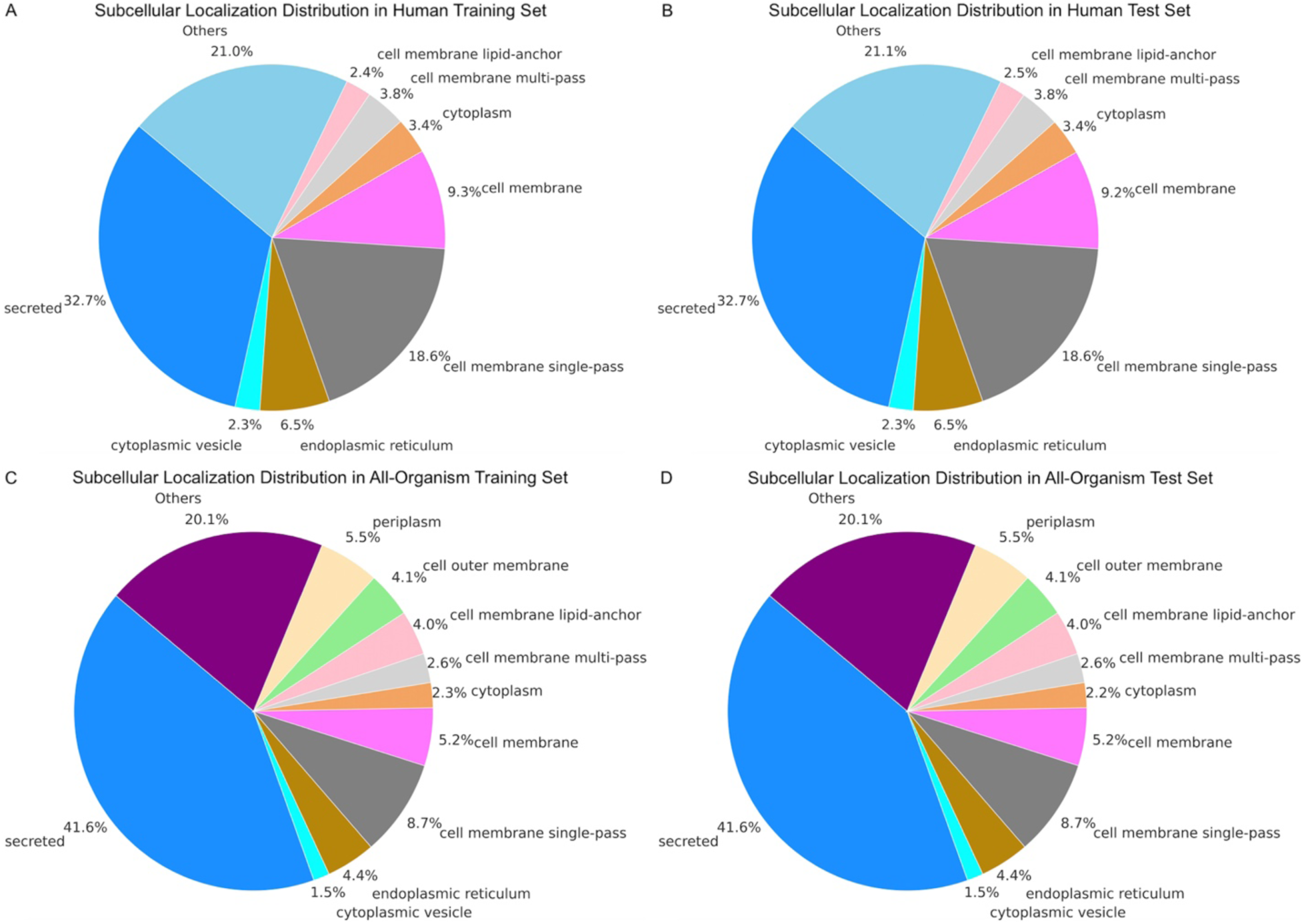
Distributions of subcellular localizations among the training and testing datasets within the human-only and all-organism datasets. **(A)** Distribution of the 11 most common subcellular locations in the human training set with the other 57 grouped under ‘Other’. **(B)** Distribution of the 11 most common subcellular localizations in the human test set with the other 57 grouped under ‘Other’. **(C)** Distribution of the 11 most common subcellular localizations in the all-organism training set with the other 57 grouped under ‘Other’. **(D)** Distribution of the 11 most common subcellular locations in the all-organism testing set with the other 57 grouped under ‘Other’.

Our ML models took in the mature protein sequences, target subcellular localizations, and organisms as inputs and generated the supposedly optimal signal peptide sequences for protein expression and localization as outputs (**Figure 2**). We incorporated an AI agent to allow for user-machine interactions in human languages. For the AI agent to start designing signal peptides, users must include the mature sequence, localization, and organism in the prompt. The AI agent is designed to inquire users to obtain all the necessary information. The information retrieved by the AI agent is then used as inputs to our ML model. The workflow, incorporating an AI agent for predicting optimal signal peptides for expressing mature proteins in a given organism with a specified cellular localization, is shown in **Figure 2**.

The performance metrics of our ML models are shown in **Figure 4**, and an example illustration of the user-software interaction with the AI agent is shown in **Figure 5**. In the next sections, we first discuss the performance of the ML model when trained on only human proteins, along with the limitations of the model and putative solutions. We then discuss the model built on proteins from all organisms and the improvements afforded by this model. Finally, we discuss the AI agent workflow and functionality.

**Figure 4.**
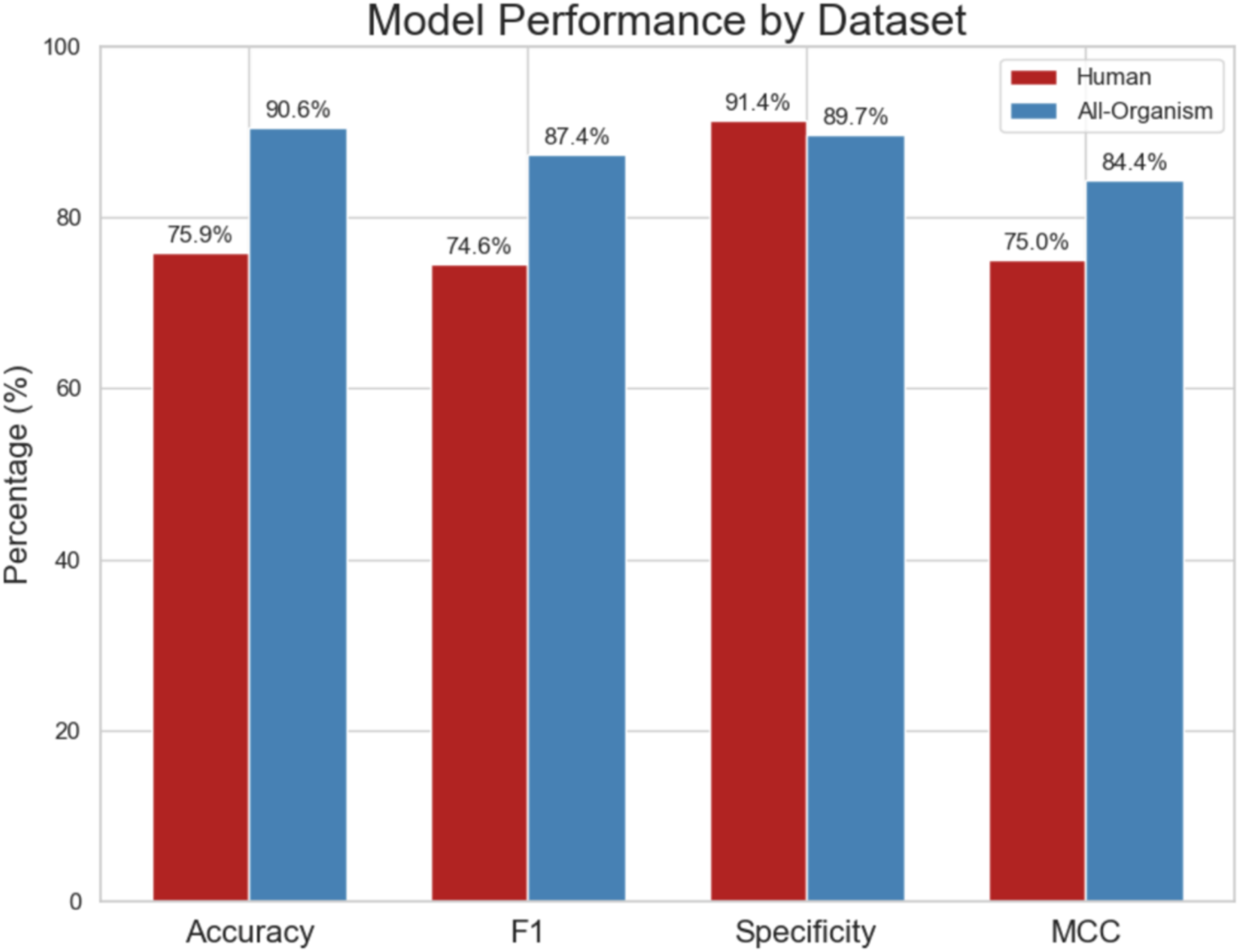
Side-by-side comparison of the performances of the ML models trained on human-only and all-organism protein data on their corresponding test sets. The plot demonstrates the superior performance of the ML model trained on all-organism protein data in terms of accuracy, F1, and MCC. The specificity is higher for the ML model trained on human-only protein data, showing prevalent class imbalances in the human-only protein dataset.

**Figure 5.**
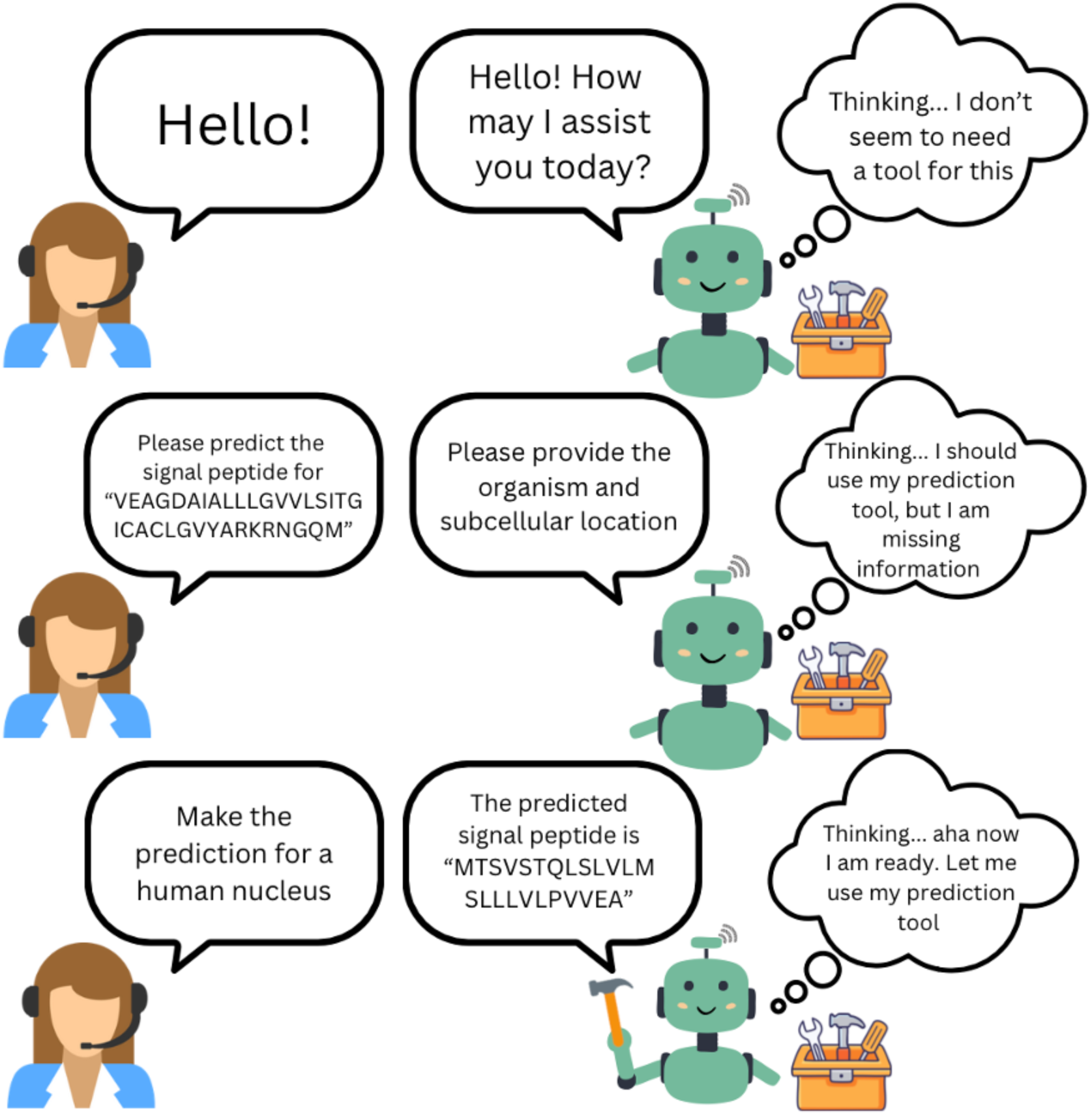
Real-world demonstration of a user interaction with the SignalGen AI agent. The incorporation of an AI agent allows for the user-software interaction through natural human languages rather than scripting languages. This figure also shows the thought process behind the SignalGen AI agent’s decisions. The AI agent passes in the inputs from the user using a custom tool to the ML model trained on all-organism protein data to make the prediction.

### Performance of the ML model to design signal peptides for optimal expression of human mature proteins, given subcellular localization in human cells

First, we trained the ML model on human-only protein data for 50 epochs. The prediction accuracy and loss of the ML model decoder on the human-only test set were logged during the training to monitor the performance of the model and is included in **Supplementary Figure 4A-B**. In particular, the accuracy of the decoder started at 19.1% and increased over the 50 epochs to 75.9% (**Supplementary Figure 4A**). During the first 20 epochs, the accuracy increased by 40.7% (**Supplementary Figure 4A**). Between epochs 20 and 50, the rate of increase in the accuracy dropped, with the accuracy only increasing by 16.1% between epoch 20 and epoch 50 (**Supplementary Figure 4A**). Meanwhile, the loss function started at 2.3 and decayed almost logarithmically before approaching a loss of 0.1 at the end of 50 epochs (**Supplementary Figure 4B**). The consistent increase in accuracy and concomitant decrease in loss shown in the plots indicated healthy and efficient training without any signs of overfitting (**Supplementary Figure 4A-4B**).

At the end of training, we evaluated the human-only ML model’s performance by testing its prediction capacity on the full sequence memory signal peptides predicted from our human-only protein test set. The confusion matrix of its predicted versus actual amino acids for each position in the human signal peptides in the test is shown in **Supplementary Figure 5A**. Based on the confusion matrix, we calculated the testing accuracy (i.e., the percentage of correct amino acids in the correct position), F1-score (i.e., the balance between precision and recall), specificity (i.e., the ability to avoid false positives), and MCC (i.e., the correlation between predicted and true token labels) for each amino acid (**Supplementary Figure 6**). The equations for these performance parameters are shown below (TP = true positive; TN = true negative; FP = false positive; FN = false negative). Here, we chose to calculate the accuracy by simply dividing the number of correct predictions over the number of predictions made to avoid inflating the metric.

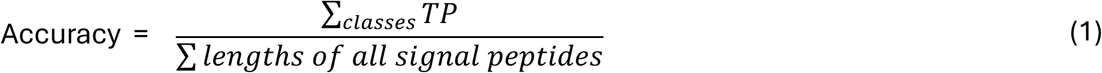

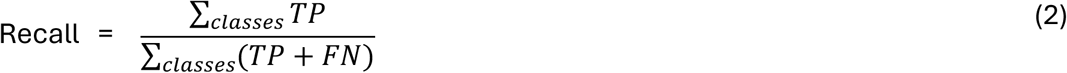

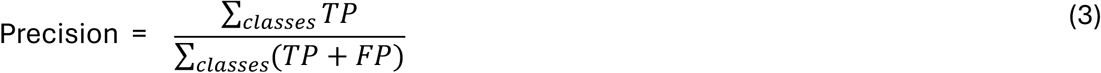

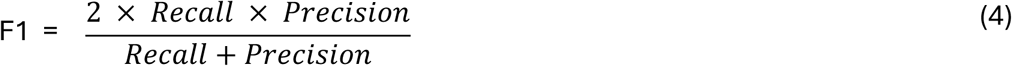

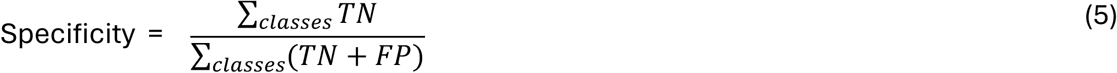

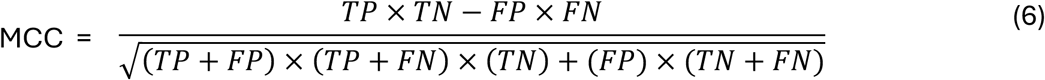

Due to the preferences of specific amino acids in the signal peptides (including leucine and alanine) (**Supplementary Figure 2**), imbalances were observed between the amino acids. Despite this, it was observed that there was little variation in the metrics between each amino acid (**Supplementary Figure 6**). This indicated that there is a rather strong balance in the predictions generated by the model. Regardless, the more common amino acids tend to have the highest values in each plot, demonstrating that the class imbalance is not completely mitigated (**Supplementary Figure 6**). We chose to calculate the micro-averaged accuracy, F1-score, specificity, and MCC and included the results in **Figure 4** to examine the performance of our human-only ML model. In particular, the accuracy was calculated to be 75.9% and the MCC was 75.0%, demonstrating the model performed relatively well in generating accurate sequences without relying on over predicting common acids (**Figure 4**). On the other hand, the F1-score was found to be 74.6%, indicating the predicted signal peptides were accurate in terms of length, while the specificity was 91.4%, showing the model was very good at avoiding false positives (**Figure 4**).

### Performance of the ML model to design signal peptides for optimal expression of mature proteins, given subcellular localizations in any organism

Next, we trained the ML model on all-organisms protein data, a larger and more diverse dataset, to further reduce class imbalance and prevent the model from overpredicting certain common amino acids in signal peptides. In fact, the dataset containing all-organisms protein data was approximately 10 times the size of the human-only dataset, with 55,620 sequences compared to 5,699 sequences (**Supplementary Data 1**). The increased size of the dataset provides more data for the model to learn from, which helps increase the generalization of the model. Furthermore, the inclusion of all organisms provides the model with the inherent ability to predict signal peptides for the mature proteins of any organism.

Here, we trained the ML model decoder on the protein data from all organisms for 50 epochs and logged the prediction accuracy as well as the loss of the ML model on the all-organism test set in **Supplementary Figure 4C-D**. After the first epoch, the ML model showed a testing accuracy of 26.8%, which increased to 90.6% by the 50^th^ epoch (**Supplementary Figure 4C**). Furthermore, the testing accuracy climbed steadily in the first 30 epochs, increasing by 55.8%, whereas in the last 20 epochs, the increase rate was much smaller, with the model accuracy only increasing by 8.0% in these last 20 epochs (**Supplementary Figure 4C**). Meanwhile, the loss value started at 3.0 after epoch one and decreased to 0.2 by the end of the 50^th^ epoch (**Supplementary Figure 4D**). In addition, the loss decreased steadily for the first 30 epochs, dropping by roughly 2.5 units, but then began to level oc during the following 20 epochs, only dropping by 0.3 units (**Supplementary Figure 4D**). Overall, the plots of accuracy and loss functions over the training courses showed that the model continuously improved throughout the 50 epochs (**Supplementary Figure 4C-D**).

At the end of training, we evaluated the performance of the all-organism ML model by testing its prediction capacity on the full sequence memory signal peptides predicted from our all-organism protein test set. The confusion matrix of its predicted versus actual amino acids for each position in the all-organism signal peptides in the test was included in **Supplementary Figure 5B**. The values within the boxes along the diagonal follow the values for the amino acid frequency for signal peptides in the dataset (**Supplementary Figure 2**), except for the value for amino acid ‘M’ being larger than the proportionate expected value. This demonstrated that the model performed well in predicting where these amino acids occur in the signal peptide sequence. Based on the confusion matrix, we calculated the testing accuracy, F1-score, specificity, and MCC for each amino acid (**Supplementary Figure 7**). We observed that these plots show the same level of variance between amino acids as the human-only model, yet the values are higher (**Supplementary Figure 7**). This indicates that a class imbalance still exists, but the model had access to a much larger dataset, enabling it to make more accurate predictions (**Supplementary Figure 7**). Due to the preferences of specific amino acids in the signal peptides (including leucine and alanine) (**Supplementary Figure 2**), imbalances were observed between the amino acids. Therefore, we again chose to calculate the micro-averaged accuracy, F1-score, specificity, and MCC and included the results in **Figure 4** to examine the performance of our all-organism ML model. We observed that the all-organism model outperforms the human-only model in each of these metrics aside from the specificity, which measures how well the model predicts true negatives (**Figure 4**). This is likely a result of the small data used in the human-only model and the consequent inability to predict signal peptides to the right length. However, if the model lacks sufficient training data to accurately predict the length of signal peptides, the number of true negatives compared to the true negatives and false positives (in accordance with formula (5)) will be high, artificially boosting the specificity.

### Incorporation of an AI agent for designing optimal signal peptides for protein expression and target localization

We incorporated an AI agent to our workflow to enhance the accessibility of our ML models to a broader userbase (**Figure 5** and **Supplementary Figure 3**). The AI agent allowed users to interact with the ML models through a natural human language rather than a computer programming language (**Figure 5**). In other words, the incorporation of an AI agent provides users with the ability to quickly utilize our ML models without the need to make code changes for their specific needs.

The AI agent, derived from GPT-4 by OpenAI, was equipped with various tools to carry out tasks related to designing signal peptides (**Figure 5**). The most relevant tool retrieved our ML model trained on all-organism protein data and provided the model with the necessary information located by the AI agent large language model (LLM) from its conversation with the users (**Figure 5**). If the AI agent LLM cannot find the necessary information in the conversation, including the mature protein, subcellular localization, and organism, it will prompt the user to provide such information (**Figure 5**). Furthermore, the AI agent is designed to store all previous interactions within the conversation and can pull all necessary information from anywhere and anytime within the conversation (**Figure 5**).

## Discussion

Despite the importance of signal peptides in protein expression and subcellular localization, there is still no effective strategy for designing optimal signal peptides for non-native or *de novo* proteins, especially considering their subcellular localizations and the expressing organisms. Current approaches for expressing proteins in human cells mainly involve randomly testing a few *ad hoc* signal peptides. In this work, we introduce SignalGen, a fully autonomous AI agent created to design optimal signal peptides for enhanced expression, regardless of whether proteins are native, non-native, or *de novo*, and target localization in specific organism cells (**Figure 2**). We trained two distinct ML models to design signal peptides based on given mature protein sequences for expression in a target organism and for directing proteins to specific subcellular localizations. Our first ML model was trained solely on human protein data from UniProt, achieving an accuracy of 75.8% after 50 epochs (**Figure 4**). Our second model was trained on protein data from all organisms in UniProt, reaching a final accuracy of 90.6% (**Figure 4**). The model trained on proteins from all organisms was integrated with an AI agent to enable users to interact with it using natural language rather than scripting, thereby increasing its accessibility (**Figure 5**).

Our ML model trained on only human proteins performed relatively well in generating signal peptides for human proteins, based on the performance metrics recorded in **Figure 4**. However, the model struggled to learn what biologically plausible signal peptides looked like for human proteins. In fact, despite being syntactically correct, the ML model trained on human-only protein data had the tendency to over-predict common amino acids (**Supplementary Figure 6**). Furthermore, the model would also repeat these amino acids sequentially, especially in the early epochs (**Supplementary Figure 8**). In other words, while it is common for certain signal peptides to have repeated amino acids, the model often over-predicts these repeated sequences (**Supplementary Figures 6** and **8**). The ML model trained on protein data from all organisms did better in learning what constitutes a biologically plausible sequence, how long the predicted signal peptide should be, and which amino acid should be predicted at a given location in the sequence. This is reflected in the superior performance of the ML model trained on protein data from all organisms rather than only humans (**Figure 4**). In particular, the ML model trained on protein data from all organisms had a higher micro-averaged specificity, F-1 score, and MCC (**Figure 4**). Furthermore, the uniformity of the metrics across amino acids illustrated the capacity of the ML model trained on all-organism protein data in predicting the correct amino acids and not predicting repetitive sequences of common amino acids (**Supplementary Figure 5B** and **7**). Given the above observations, our ML model trained on all-organism protein data should be ready for applications to design signal peptides for expressions of both existing and *de novo* proteins.

After developing our ML model, we incorporated an AI agent, derived from the OpenAI AI agent, into SignalGen to allow for user-software interactions using natural human languages rather than scripting languages (**Figure 5**). In its current state, SignalGen incorporates our ML model trained on all-organism protein data and several other tools that enable it to dynamically handle and adjust to various situations, including different file types, dataset formats, and prompts (**Figures 2, 5** and **Supplementary Figure 3**). In addition, SignalGen can guide users through the prediction workflow if users need help using the software. The incorporation of various tools in SignalGen helps enhance user-friendliness, especially for users with limited background knowledge of the model or what constitutes a valid prompt. Overall, SignalGen represents an advancement in integrating AI agents with biological tools, thereby strengthening the connection between humans and machines (**Figures 2, 5** and **Supplementary Figure 3**).

While multiple ML-based signal peptide prediction tools have been developed and some of which also utilize the protein language model, for example, ProtBert^22^, no prediction tool has been developed to incorporate the organism and subcellular localization of the protein in designing signal peptides for optimal protein expression^10–13^. SignalGen incorporated a unique framework to take into account the mature protein sequences, target subcellular localizations, and expressing organisms in designing the optimal signal peptides (**Figures 2, 5** and **Supplementary Figure 3**). Since our model is unique, it is not feasible to make direct comparisons of its performance to the others currently available. In particular, SecretoGen exclusively generates signal peptides for protein secretion, whereas SignalGen generates signal peptides for target subcellular localization anywhere inside or outside the cell^12^. Therefore, we will not attempt to categorize these tools as the same, nor directly compare their performances. Consequently, we can only benchmark our models using well-recognized ML metrics (including accuracy, F1-score, specificity, and MCC) and compare performances between the two models trained for this study.

For future studies, we will continuously fine-tune our trained ML models as more protein data become available on the UniProt database (https://www.uniprot.org/uniprotkb). We expect the model performance to become even better as we can train the ML models on more than 5,699 human protein data and 55,620 all-organism protein data available in UniProt (https://www.uniprot.org/uniprotkb) as of July 2025. Our expectation is validated, given the notable performance increase from the ML model trained on 5,699 human sequences to 55,620 all-organism sequences (**Figure 4**). Furthermore, we may consider a custom physics-informed loss function^23^ and consider structural features of the mature proteins in training the ML models. Lastly, we may incorporate proteins with no signal peptides in our training set and train our model in a semi-supervised manner^24,25^. We expect these directions to further improve the performance of our ML models.

In summary, we developed SignalGen, an AI-agent incorporated ML model that effectively designed signal peptides for optimal protein expression of non-native and *de novo* proteins in given organisms and target subcellular localizations. SignalGen will be especially valuable for researchers who design de novo proteins, as it can be used to increase secretion in a specific subcellular compartment for a given organism. Lastly, SignalGen will be a valuable tool for the mRNA vaccine platform^5,26^ when viral or bacterial mRNA for antigens needs to be expressed in human cells.

## Methods

### Collection of protein datasets for training and evaluating SignalGen

First, we collected 3,568 reviewed human protein sequences from the UniProt Knowledgebase (https://www.uniprot.org/uniprotkb/) as of July 2025. Each protein was required to include an experimentally verified subcellular location and signal peptide sequence. The sequences obtained from UniProt were in their immature forms, which were then split into the signal peptide and mature sequences. Furthermore, it should be noted that most of these proteins had multiple subcellular localizations. Therefore, we chose to treat each different subcellular localization as a separate entry for training our ML models. After separating these entries and cleaning the data of any duplicates, there was a total of 5,699 human-only data points for training our ML model (**Supplementary Data 1**). Among these proteins, the majority had signal peptides of 10 and 40 amino acids in length (**Supplementary Figure 1A**). However, several signal peptides of 3 to 10 amino acids long were observed among the dataset (**Supplementary Figure 1A**). On the mature protein sequence side, most of the mature protein sequences were between 0 and 2,000 amino acids in length (**Supplementary Figure 1B**). Within the signal peptide sequences, there were some amino acids that were more common than others, such as A, G, L, and S (**Supplementary Figure 2A**).

The dataset consisting of human-only protein data was split into training and testing sets so that the performance of our ML model can be monitored during training. In particular, the training set was comprised of 80% of the total dataset, while the testing dataset comprised of the other 20% (**Supplementary Data 1**). The training and test sets composed of randomly sampled entries from the full dataset to introduce variation and diversify the sets. Seemingly, the split resulted in no overlap between the training and testing sets (**Supplementary Data 1**). Additionally, to ensure that the model was trained on every subcellular location and a representative number of entries from each subcellular localization, the datasets were stratified based on the subcellular localizations (**Supplementary Data 1**). We observed that the training and testing datasets maintained similar distributions of subcellular localizations (**Figure 3**). The proteins in the training and test sets were randomly selected from the original dataset based on the number of entries contained in each subcellular localization. In the human-only protein dataset, we found 68 different subcellular localizations, with the most common localization being ‘Secreted’ (accounting for 32.7%) and the other 57 not labeled falling under the category ‘Other’ (accounting for 21.0%) (**Figure 1A**).

Later, we collected the protein data for all organisms in a similar manner. The total number of proteins from all organisms with experimentally validated signal peptides and subcellular localizations was 40,610 from UniProt as of July 2025. After following the described processes of separation and removal of duplicates, the dataset contained 55,620 all-organism data points for training the ML models (**Supplementary Data 1**). The entries from all-organism proteins showed similar distributions in terms of the lengths of signal peptides (**Supplementary Figure 1C**), lengths of mature proteins (**Supplementary Figure 1D**), and amino acid compositions in signal peptides to the human proteins (**Supplementary Figure 2B**). However, we observed more outliers in the lengths of all-organism signal peptides, with some sequences up to 140 amino acids long (**Supplementary Figure 1C**). For the all-organism dataset, we also only stratified the dataset on the subcellular localizations rather than both the subcellular localizations and organisms because over 2,000 organisms out of the over 4,000 different organisms only appeared once in the dataset. Consequently, the number of different groups to stratify the data upon for both organisms and subcellular localizations would not have been computationally efficient. Here, we also randomly split the full dataset into the training and test sets using an 80:20 ratio (**Supplementary Data 1**). The training and test sets for the all-organism ML model again showed similar distributions to each other in terms of subcellular localizations (**Figure 3**).

### Training protocol for SignalGen

Our ML models took in the amino acid sequences of the mature proteins, their subcellular localization, and the names of their organisms as inputs (**Figure 2**). The final output was the predicted amino acid sequence of the signal peptides for optimal expression of the mature proteins in the given subcellular localizations and organisms. Our ML models utilized a Latent Residual Transformer framework^27^. The amino acid sequences of the mature proteins were embedded using the Seq2Seq transformer model^28^. The Seq2Seq transformer model effectively takes in a vector of characters and output a vector of characters while understanding patterns within the sequences^29^. Here, we used ProtBert^22,30,31^, a protein language model trained on protein sequences, to encode the amino acid sequences. ProtBert was pretrained using approximately 216 million proteins from UniRef100 and uses learned embeddings to capture the structure and qualities of the protein within the numerical vector^30^. The subcellular localizations and organism names were embedded as labels.

We defined the vocabulary for the amino acid sequences which included all canonical amino acid as well <START>, <END>, and <PAD>. Here, <START>, <END>, and <PAD> were tokens used for starting, ending, and filling the predictions of amino acids for signal peptides. We also defined the vocabulary for the two labels using numerical mapping. The majority of the mature protein amino acid sequences had lengths under 2,000 amino acids. However, to use ProtBert to encode an embedding, each sequence had to be under 1,024 amino acids long^22^. To train a more versatile model without compromising computing efforts, we decided to use the maximum sequence length of 1,024 amino acids (**Supplementary Figure 1**). As a result, any mature protein sequences under the maximum value were padded with the pad value, and any sequences over the maximum value were truncated by removing the amino acids past the maximum value index.

The mature sequences and signal peptides were tokenized using an AutoTokenizer^27^ that would format each input to the ProtBert^22^ specifications by adding spaces between amino acids. To train the decoder and teach it to generate sequence of the correct length, <START> was prepended while <END> was appended based on Bahdanau et al^32^. We encoded the signal peptide sequences into full sequence memory and latent sets. The labels were tokenized using the factorization function from the Pandas library^33^. The mature sequences were then passed through ProtBert^22^ using the same AutoTokenizer as used for the signal peptides, which returned the sequence as a vector where each token helped represent the protein. The mature protein was only encoded into a latent set, and it was aligned with the signal peptide using contrastive learning. The signal peptide full sequence memory and latent sets were encoded using cross-attention with the mature protein vector, localization, and organism in the case of the all-organism model^32^ (**Supplementary Figure 3**). Each of the labels got their own auxiliary head and was embedded using multi-head attention pooling; thus, the model was able to learn different patterns and relationships between the mature protein sequences, each of the labels, and the optimal signal peptide sequences (**Figure 2** and **Supplementary Figure 3**). The usage of a multi-head attention pooling is especially important for designing signal peptides because each of the labels has an important role in the function of the signal peptide^1^. Notably, we used an auxiliary stop head for the end of the sequence to help the model predict when to stop the sequence. The usage of an auxiliary stop head was an improvement upon using <END> as a special token because the model would treat it similar to any other amino acid rather than an indication to stop predicting amino acids. After the mature sequence latent set is mapped to the signal peptide latent set, it is refined using the two residual transformers in the LRT model. The resulting predicted signal peptide latent set is expanded into a full sequence memory representation with slight noise and inputted to the decoder.

The dataset of human and all-organism proteins from UniProt showed notable imbalances across the amino acids due to the preferences of certain amino acids in signal peptides^5^ (**Supplementary Figure 2**). To prevent the decoder from overpredicting the common amino acids in signal peptides to artificially raise the accuracy, we used a focal loss function^34^ that forces the model to take risks and predict rarer amino acids. Part of this implementation included adding class weights that penalized overpredicting common amino acids^35^. In addition, the loss function was modified to ignore the padding appended to each vector. Furthermore, we added an auxiliary loss function that uses a discriminator to differentiate between plausible and implausible signal peptide sequences to ensure that the loss was helping the ML models predict biologically plausible as well as mitigating the effects of class imbalance in **Supplementary Figure 2**. We combined the evaluation of plausibility from the auxiliary loss function with the focal loss. Notably, we also created and added a loss function that would compute a biological plausibility score based on the hydrophobicity and charge of the predicted signal peptide compared to the true signal peptide. For each prediction, we also used a domain aware scoring-based reward function we call Fast In Silico Sequence Selection and Triage (FISST) similar to the method outlined in Ranzato et al^36^. This extracted the n-region and h-region from each prediction and rewarded the model based on the accuracy of the hydrophobicity and charge to further reinforce biological plausibility of the predicted signal peptides.

We used TransformerDecoder, a decoder from PyTorch^37^, to generate the signal peptide amino acid sequence from the signal peptide full sequence memory representation. The decoder was based on the methods introduced by Raffel et al^38^. The decoder was applied with a mask to keep the model from looking ahead. During inference, the model started from the <START> token and predicted one amino acid at a time until the model reached the auxiliary stop head. The ML models employed the optimizer of AdamW^39^ with a learning rate of 1×10^-4^. Due to the complexity of the all-organism dataset with over 40,000 entries and over 4,000 different organisms, the ML model trained on all-organism protein data was trained using slightly different parameters than the ML model trained on human-only protein data. To keep the model from being overwhelmed in early training and crashing, we used a higher value for label smoothing as well as a more gradual decay of the α value in our focal loss function.

The SignalGen AI agent was designed using LangChain^40^. LangChain is a framework for designing large language model (LLM)-based applications^40^. This allowed for the creation of a waterfall architecture where a main agent can delegate to different subagents depending on the prompt and task, as outlined in Zhang et al^41^. In turn, each of these subagents can delegate to their own subagents. The delegations enable the efficient transmission of information from prompts into tools and agents that can quickly decipher which tools to use. In the case of SignalGen, the main agent has three possible paths, including delegating to a subagent to check for the necessary information to use the ML model predicting signal peptides for mature proteins across all organisms, delegating to a subagent to perform file related operations, or responding to prompts without using any prediction tools. Each agent was based on the OpenAI ChatGPT LLM, allowing the agents to parse natural languages and respond in natural languages. ChatGPT was not used for anything other than facilitating natural language parsing and responses.

## Data Availability Statement

Data supporting the findings of this study are included in the article and its Supplementary Information files.

## Code Availability Statement

The custom codes related to SignalGen are related to a provisional patent application that was filed by Los Alamos National Laboratory (LANL) Richard P. Feynmann Center for Innovation on November 11^th^, 2025.

## Conflict of Interest Statement

The authors declare no conflicts of interest.

## Acknowledgements

This work was supported by the Defense Threat Reduction Agency under the Rapid Assessment of Platform Technologies to Expedite Response (RAPTER) program. The views expressed in this article are those of the authors and do not reflect the official policy or position of the U.S. Department of Defense or the U.S. Government. Our ML models, along with the associated scripts and data generated by the models, will be made available upon proper clearance from our funding sponsors. We would like to thank Los Alamos National Laboratory Institutional Computing for their computing resources. The authors thank Dr. Traci Pals and Dr. Bob Webb for their support of this work.

## Author Contributions

J.K. performed research, analyzed data, and wrote manuscript. H.N.D. performed and supervised research, analyzed data, and wrote manuscript. A.S. wrote manuscript. J.Z.K. acquired funding, supervised research, and wrote manuscript. S.G. supervised research, analyzed data, and wrote manuscript. All authors contributed to the final version of the manuscript.

## Notes

### Competing Interest Statement

J.H.M., H.N.D., J.Z.K., and S.G. are inventors of a provisional patent application submitted by Los Alamos National Laboratory (LANL) Richard P. Feynman Center for Innovation for the development of SignalGen on November 11th, 2025.

