## Supplementary Information for "SignalGen: A Protein Language Model Based AI Agent For Optimal Signal Peptide Prediction"

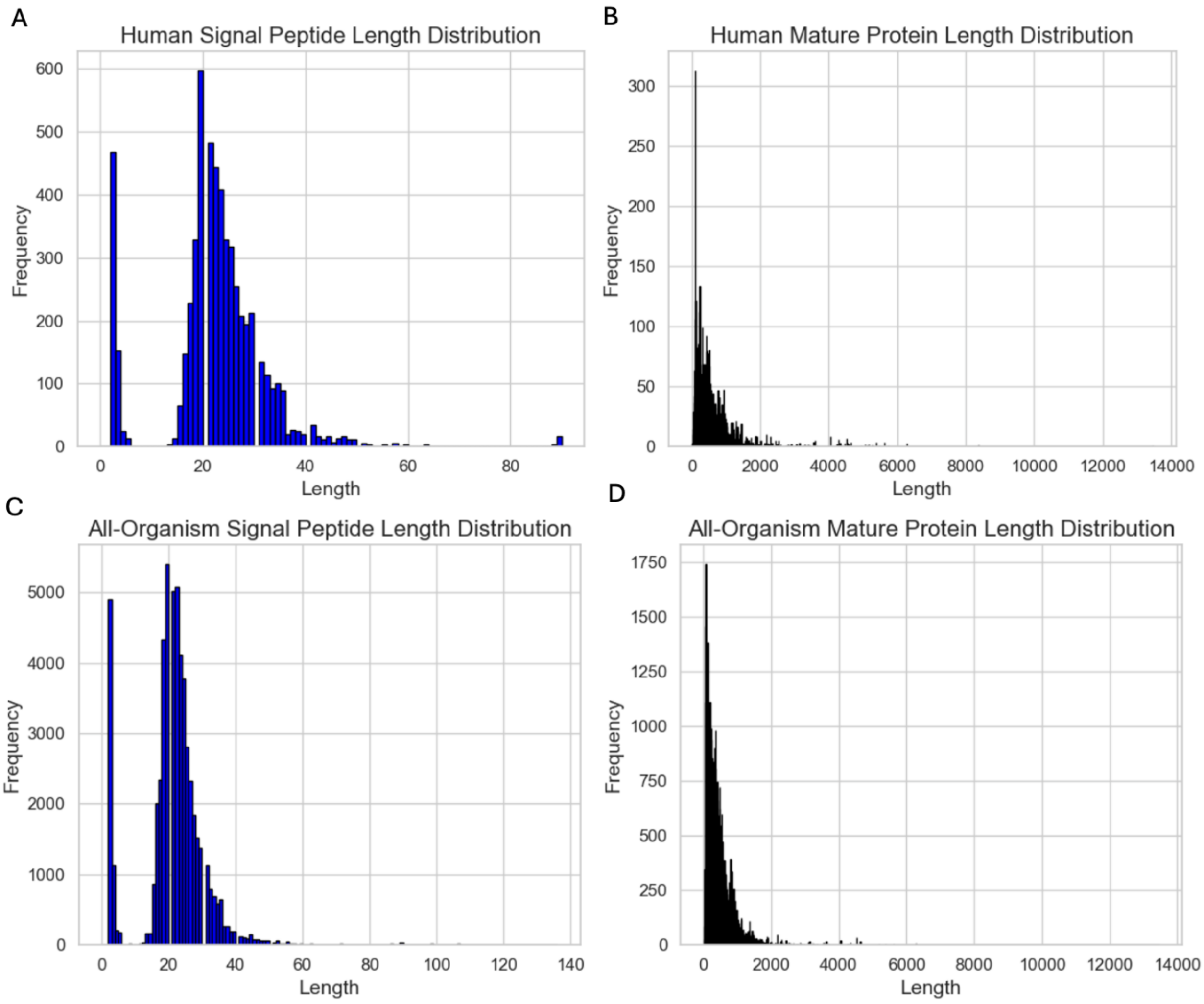


**Supplementary Figure 1. Distributions of the sequence lengths of the signal peptides and mature proteins in human-only and all-organism datasets. (A)** Frequency of different sequence lengths observed for human signal peptides. **(B)** Frequency of different sequence lengths observed for human mature proteins. **(C)** Frequency of different sequence lengths observed for all-organism signal peptides. **(D)** Frequency of different sequence lengths observed for all-organism mature proteins.

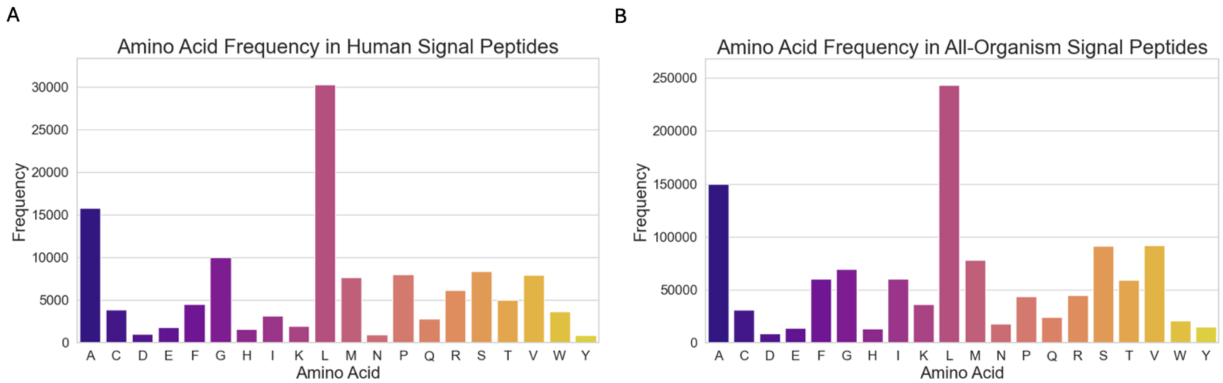


**Supplementary Figure 2. Distributions of amino acid frequencies observed for human and all-organism signal peptides.** **(A)** Frequency of each amino acid appearance within human signal peptides. **(B)** Frequency of each amino acid appearance within all-organism signal peptides.

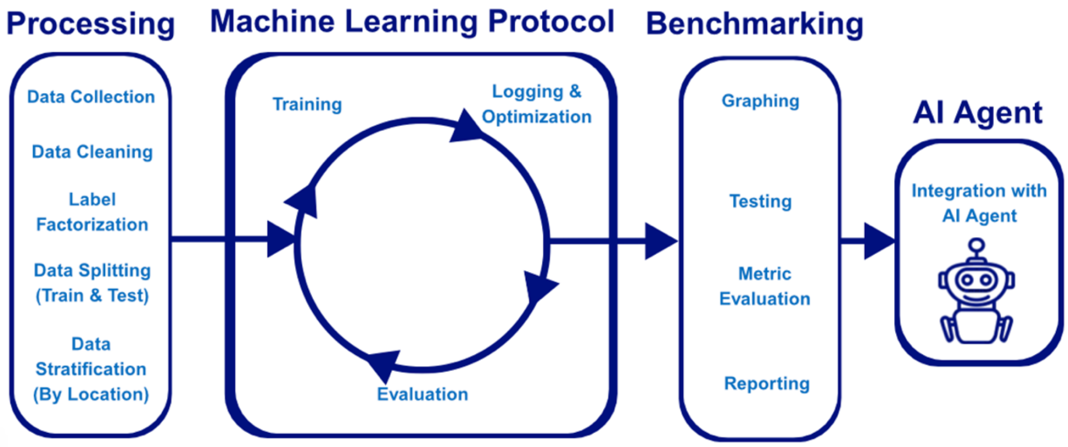


**Supplementary Figure 3. End-to-end project workflow from data processing to incorporation of an AI agent.** The figure demonstrates each step taken in the three key steps to developing the ML models: processing, ML protocol, and benchmarking. At the fourth step, we included an AI agent to enhance user accessibility.

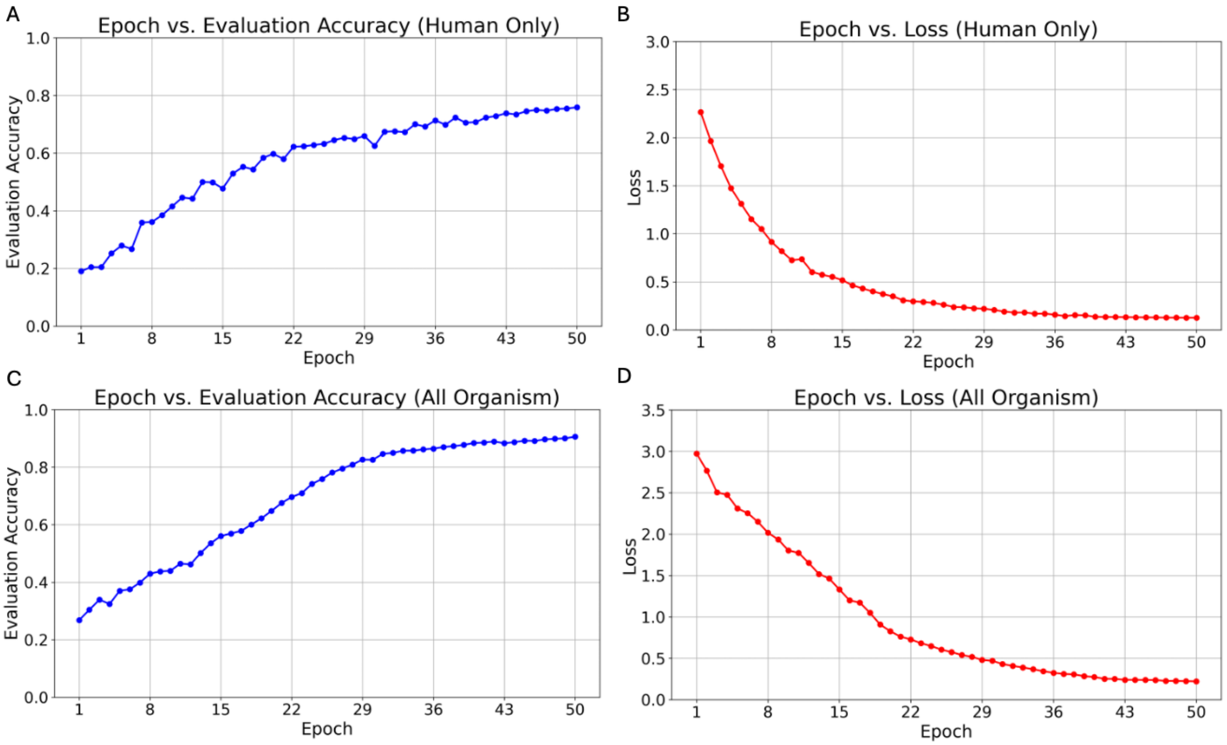


**Supplementary Figure 4. Plots of the testing accuracy and loss functions observed during training for the human and all-organism models. (A)** Accuracy of the ML model trained on all-organism protein data for 50 epochs. **(B)** Loss of the ML model trained on all-organism protein for 50 epochs. **(C)** Accuracy of the ML model trained on human-only protein data for 50 epochs. **(D)** Loss of the ML model trained on human-only protein data for 50 epochs.

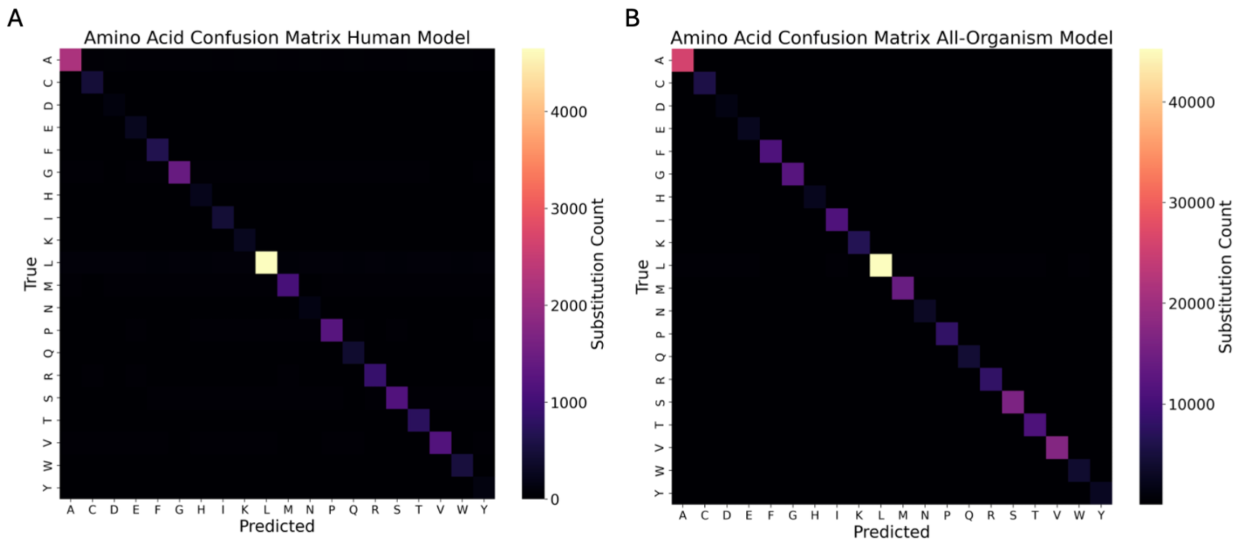


**Supplementary Figure 5. Confusion matrices of predicted versus real amino acids in the human and all-organism signal peptides made by SignalGen on the respective test sets.** **(A)** Confusion matrix for the predictions made by the ML model trained on human-only protein data. **(B)** Confusion matrix for the predictions made by the ML model trained on all-organism protein data.

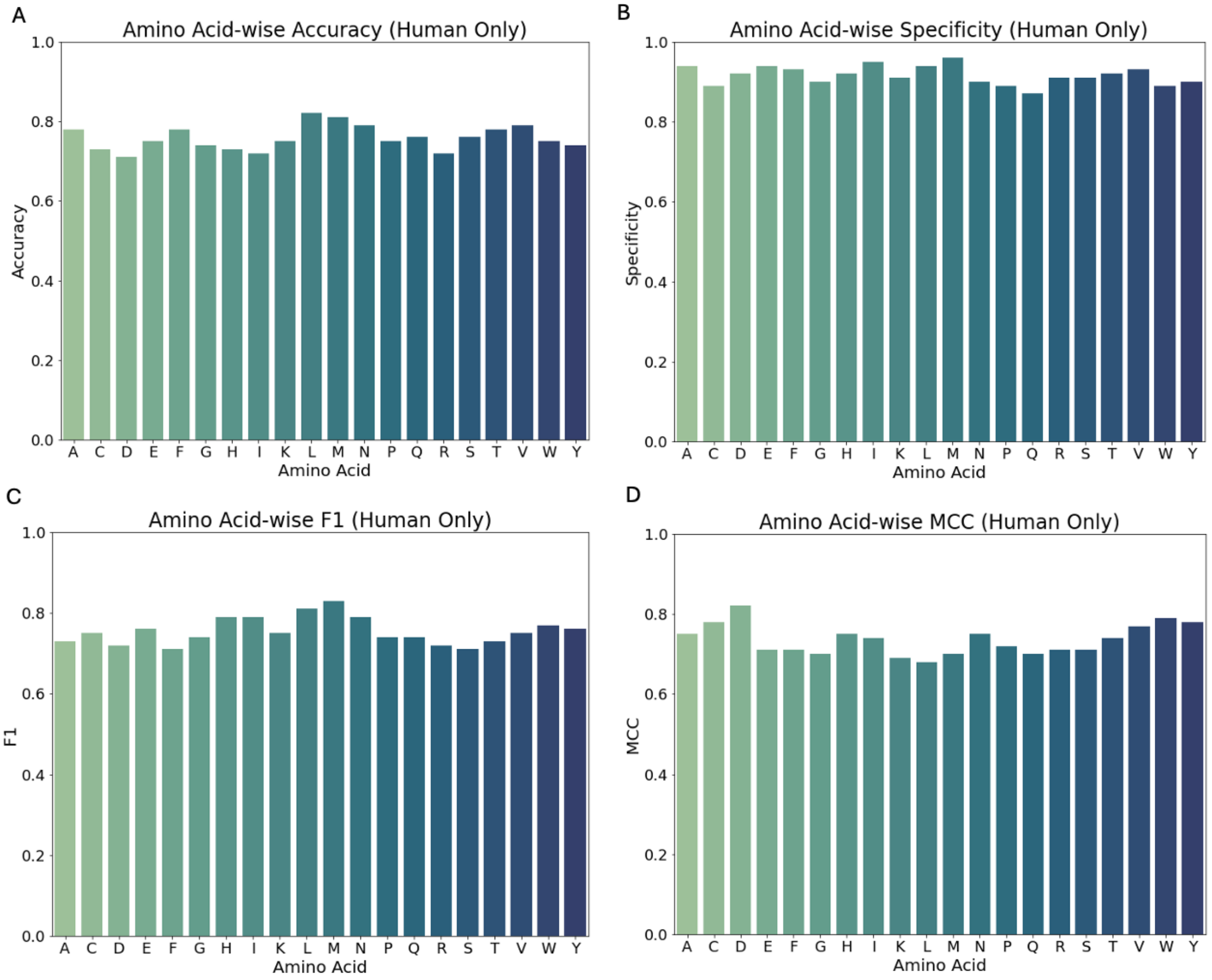


**Supplementary Figure 6. Performance metrics across amino acids for predictions made by the ML model trained on human-only protein data. (A)** Amino acid-wise accuracy of the ML model trained on human-only protein data. **(B)** Amino acid-wise specificity of the ML model trained on human-only protein data. **(C)** Amino acid-wise F1-score of the ML model trained on human-only protein data. **(D)** Amino acid-wise MCC of the ML model trained on human-only protein data.

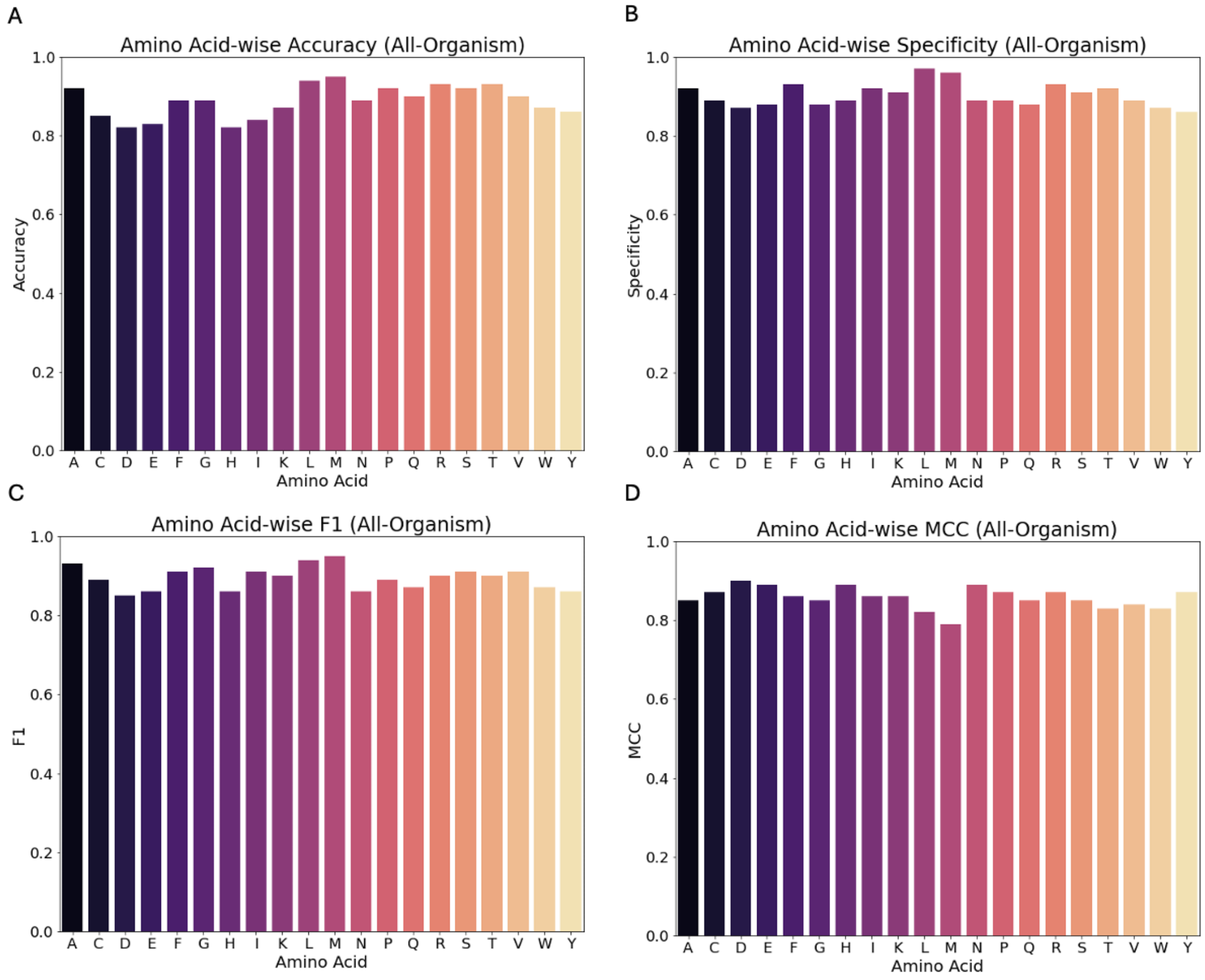


**Supplementary Figure 7. Performance metrics across amino acids for predictions made by the ML model trained on all-organism protein data. (A)** Amino acid-wise accuracy of the ML model trained on all-organism protein data. **(B)** Amino acid-wise specificity of the ML model trained on all-organism protein data. **(C)** Amino acid-wise F1-score of the ML model trained on all-organism protein data. **(D)** Amino acid-wise MCC of the ML model trained on all-organism protein data.

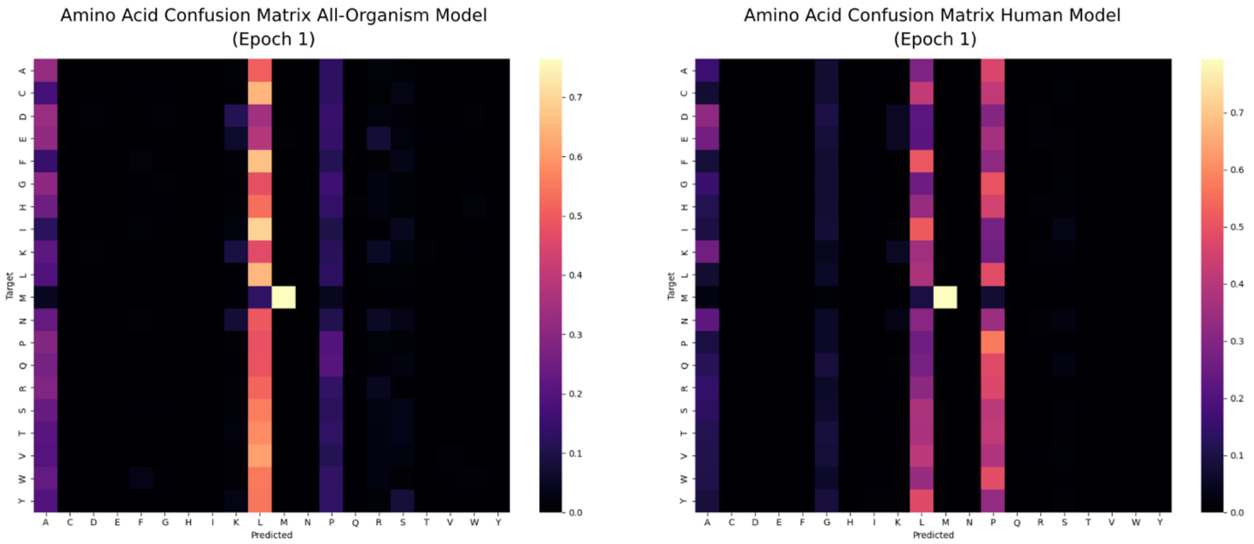


**Supplementary Figure 8. Amino acid confusion matrices for the human and all-organism models after epoch 1. (A)** The amino acid confusion matrix for the all-organism model after being trained for 1 epoch. **(B)** The amino acid confusion matrix for the human model after being trained for 1 epoch.
